# Multiple infiltration and cross-species transmission of foamy viruses across Paleozoic to Cenozoic era

**DOI:** 10.1101/2020.12.18.423569

**Authors:** Yicong Chen, Yu-Yi Zhang, Xiaoman Wei, Jie Cui

## Abstract

Foamy viruses (FVs) are complex retroviruses that can infect humans and other animals. In this study, by integrating transcriptomic and genomic data, we discovered 412 FVs from 6 lineages in amphibians, which significantly increased the known set of FVs in amphibians. Among these lineages, salamander FVs maintained a co-evolutionary pattern with their hosts that could be dated back to the Paleozoic era, while, on the contrary, frog FVs were much more likely acquired from cross-species (class level) transmission in the Cenozoic era. In addition, we found three distinct FV lineages had integrated into the genome of a salamander. Unexpectedly, we identified a potential exogenous form of FV circulated in caecilian, demonstrating the existence of exogenous form of FV besides mammals. Our discovery of rare phenomena in amphibian FVs has overturned our collective understanding of the macroevolution of the complex retrovirus.

**Importance:** Foamy viruses (FVs) represent, more so than other viruses, the best model of co-evolution between a virus and a host. This study represents so far, the largest investigation of amphibian FVs and revealed 412 FVs of 6 distinct lineages from three major orders of amphibians. Besides co-evolutionary pattern, cross-species and repeated infection were also observed during evolution of amphibian FVs. Remarkably, expressed FVs including a potential exogenous form were discovered, suggesting live FVs could be underestimated in nature. These findings revealed the multiple origin and complex evolution of amphibian FVs started from the Paleozoic era.

## Introduction

Retroviruses (family *Retroviridae*) have great medical and economic significance, as some are associated with severe infectious disease or are oncogenic(1-5). Retroviruses are notable as they occasionally integrate into the germline of a host and become endogenous retroviruses (ERVs), which can be vertically inherited(6, 7). ERVs generated by simple retroviruses are widely distributed in vertebrates(8-17). However, complex retroviruses, such as lenti-, foamy- and delta-retroviruses, have rarely appeared as endogenous forms.

Foamy viruses (FVs) (subfamily *Spumavirinae*) are complex retroviruses that exhibit typical co-divergence with their host, providing an ideal framework for understand the long-term evolutionary relationship between viruses and vertebrate hosts(18-21). Exogenous foamy viruses are prevalent in mammals, including primates(18, 22), bovines(23, 24), equines(25), bats(26), and felines(27). Vertebrate genomic analysis first identified endogenous foamy viruses (EFVs) in sloths(28), then they were found in several primate genomes(29-31). The subsequent discovery of EFVs and EFV-like elements in fish and amphibian genomes indicated that foamy viruses, along with their vertebrate hosts, have ancient origins(32, 33). Recently, the discovery of four novel reptile EFVs and two avian EFVs demonstrated that FVs can infect all five major classes of vertebrates(34-37).

Although substantial number of EFVs had been found across the evolutionary history of vertebrates(28, 30, 38), to date, only a few of them have been found in amphibians, mainly in salamander(32). Most importantly, no potential exogenous foamy viruses have been found outside of mammalian host(32, 37). In this study, by integrating transcriptomic and genomic data, we found >400 FVs of 6 novel lineages of foamy viruses, which significantly increased the set of known foamy viruses in amphibians. Several expressed FVs were revealed, with potential exogenous foamy viruses discovered. Our work also provides novel insight into the early evolution of foamy viruses.

## Results

### Discovery and confirmation of foamy viral elements and expressed foamy viruses in amphibians

We screen 19 amphibian genomes (Table S1) using a homologous-based stepwise manner as described in the materials and methods section. This led to the discovery of 18 foamy-like ERVs in *Spea multiplicate* (Mexican spadefoot toad), 64 in *Rhinatrema bivittatum* (two-lined caecilian), and 131 in *Ambystoma mexicanum* (axolotl) (Table S2). The foamy-like elements in *S. multiplicate* showed high similarity with each other. Such high similarity between copies has also been observed in foamy-like elements in *R. bivittatum*. However, there were two distinct groups of foamy-like ERVs in *A. mexicanum* that showed an average 64% identity with 99% coverage.

In order to discover any potential exogenous foamy viruses, we also searched all available amphibian RNA-seq data in the transcriptome sequence assembly (TSA) database. Notably, we also found 58 foamy-like RNA copies in *Tylototriton wenxianensis* (Wenxian knobby newt), 131 in *Taricha granulosa* (Rough-skinned Newt), and 10 in *R. bivittatum*. Homologous comparison with NviFLERV-1 showed that all three groups of foamy-like viral copies harbored typical major coding regions, i.e., GAG, POL and ENV, with an average similarity of 22%–76%.

To examine whether these viral elements belonged to the clade of foamy viruses, a phylogenetic tree was inferred using the reverse transcriptase (RT) protein of representative viruses from all of the genera of *Retroviridae* (Fig. 1 and Table S3). Our RT phylogenetic tree revealed that novel viral elements found in amphibians were grouped within the foamy virus clade (bootstrap value > 90), which confirmed that they were indeed foamy viruses. In addition, these foamy-like viral elements could be divided into three clades, where the viral elements found in different newts, including previously identified NviFLERV, clustered together, while the viral elements found in *R. bivittatum* and *S*.*multiplicate* both formed a monophyletic group, respectively. Also, of note was that foamy-like ERVs in *A. mexicanum* divided into two clusters, indicating the existence of two lineages of EFVs in *A. mexicanum*.

**Fig. 1.**
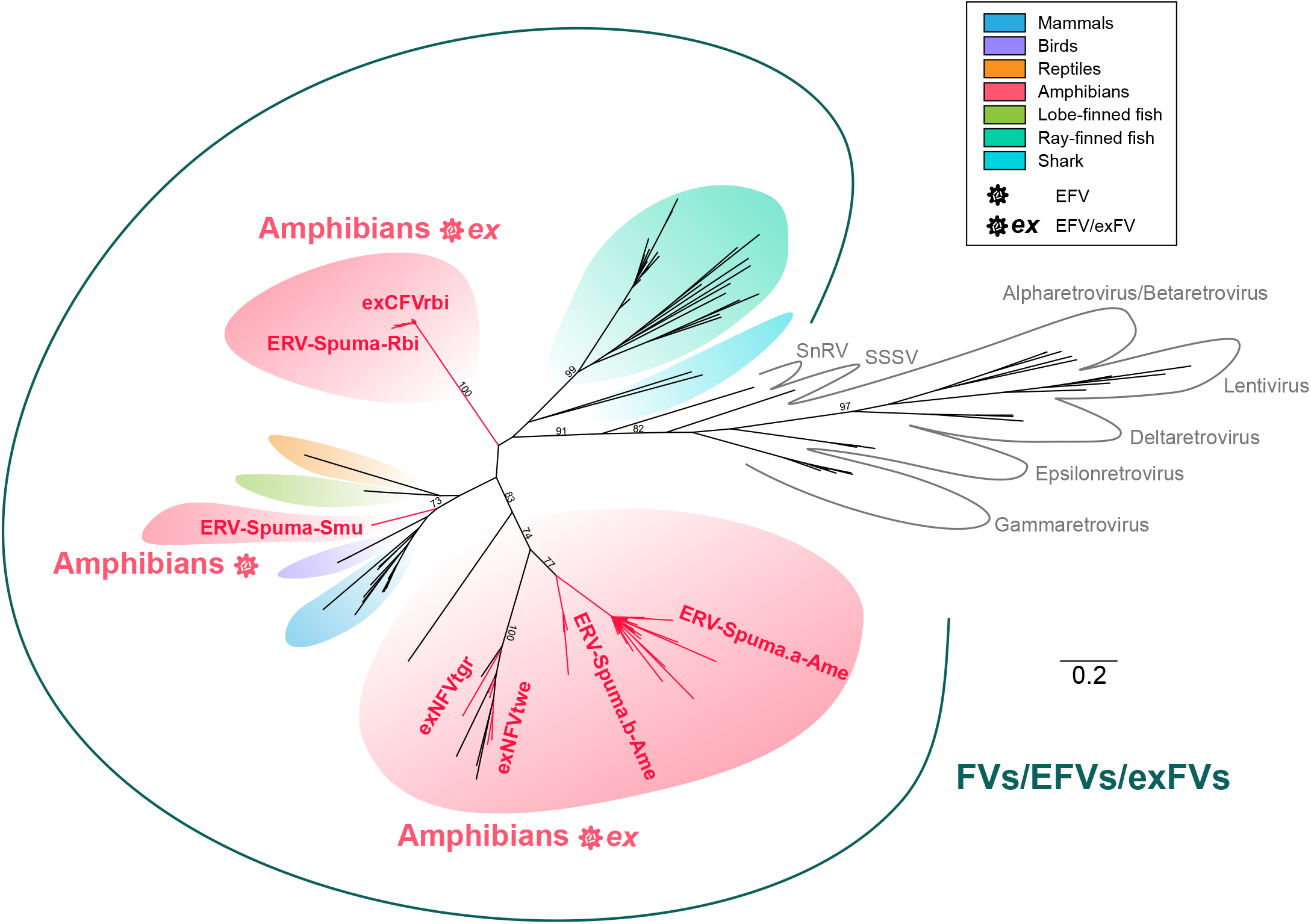
Unrooted phylogeny of retroviruses and foamy-like elements. The tree was inferred from reverse transcriptase (RT) protein alignment. The host information of FVs, EFVs and exFVs is indicated using shaded box. The newly identified viral elements are labeled in red. The scale bar indicates the number of amino acid changes per site. Bootstrap values <70 percent are not shown.

Thus, in accordance with the nomenclature proposed for ERVs(39), the foamy-like ERVs found in *S. multiplicate* and *R. bivittatum* were named ERVs-Spuma.n-Smu (n = 1∼18) and ERVs-Spuma.n-Rbi (n = 1∼64), respectively. Additionally, two lineages in *A. mexicanum* were designated as ERVs-Spuma.an-Ame (n = 1∼104) and ERVs-Spuma.bn-Ame (n = 1∼18), respectively. Moreover, the expressed viral elements found in *T. wenxianensis, T. granulosa* and *R. bivittatum* were then named expressed NFVtwe.n (exNFVtwe) (n = 1∼58), expressed NFVtgr.n (exNFVtgr) (n = 1∼131) and expressed CFVrbi.n (exCFVrbi) (n = 1∼10), respectively.

### EFV and exFV genome characterization

14 ERVs-Spuma-Smu, 54 ERVs-Spuma-Rbi, 57 ERVs-Spuma.a-Ame and 15 ERVs-Spuma.b-Ame, which were long (> 5 kb in length) were retrieved from their respective genomes, and they were used to construct a consensus sequences for each EFV lineage (Fig. 2A and Data set S1). The consensus genomes of novel EFVs all harbored long pairwise LTRs and exhibited typical foamy virus structure, containing three major genes, GAG, POL, and ENV, and one (ERV-Spuma-Rbi) or two (ERVs-Spuma-Smu, ERVs-Spuma.a-Ame and ERVs-Spuma.b-Ame) putative accessory genes. It is noteworthy that these accessory genes all showed no similarity to any known proteins. By searching against the Conserved Domain Database (CDD) using CD-Search, we were able to identify two typical foamy virus conserved domains in all four consensus EFVs: (1) the Gag_spuma super family domain (cl26624)(40, 41) and the (2) Foamy_virus_ENV super family domain (cl04051)(38). However, ERV-Spuma-Smu exclusively contained the Spuma_A9PTase super family domain (cl08397)(32) that is present in all mammalian foamy virus Pol proteins. The existence of these domains gave additional support to their classification as foamy viruses. Other regions or domains, such as the RT_like super family (RT) domain (cl02808), the RNase_H_like super family and RT_RNaseH_2 super family (RH) domains (cl14782, cl39038), and the rev and Integrase_H2C2 and SH3_11 super family domains (INT) (pfam00665, pfam17921, cl39492) were also identified in all EFVs (Fig. 2A).

**Fig. 2.**
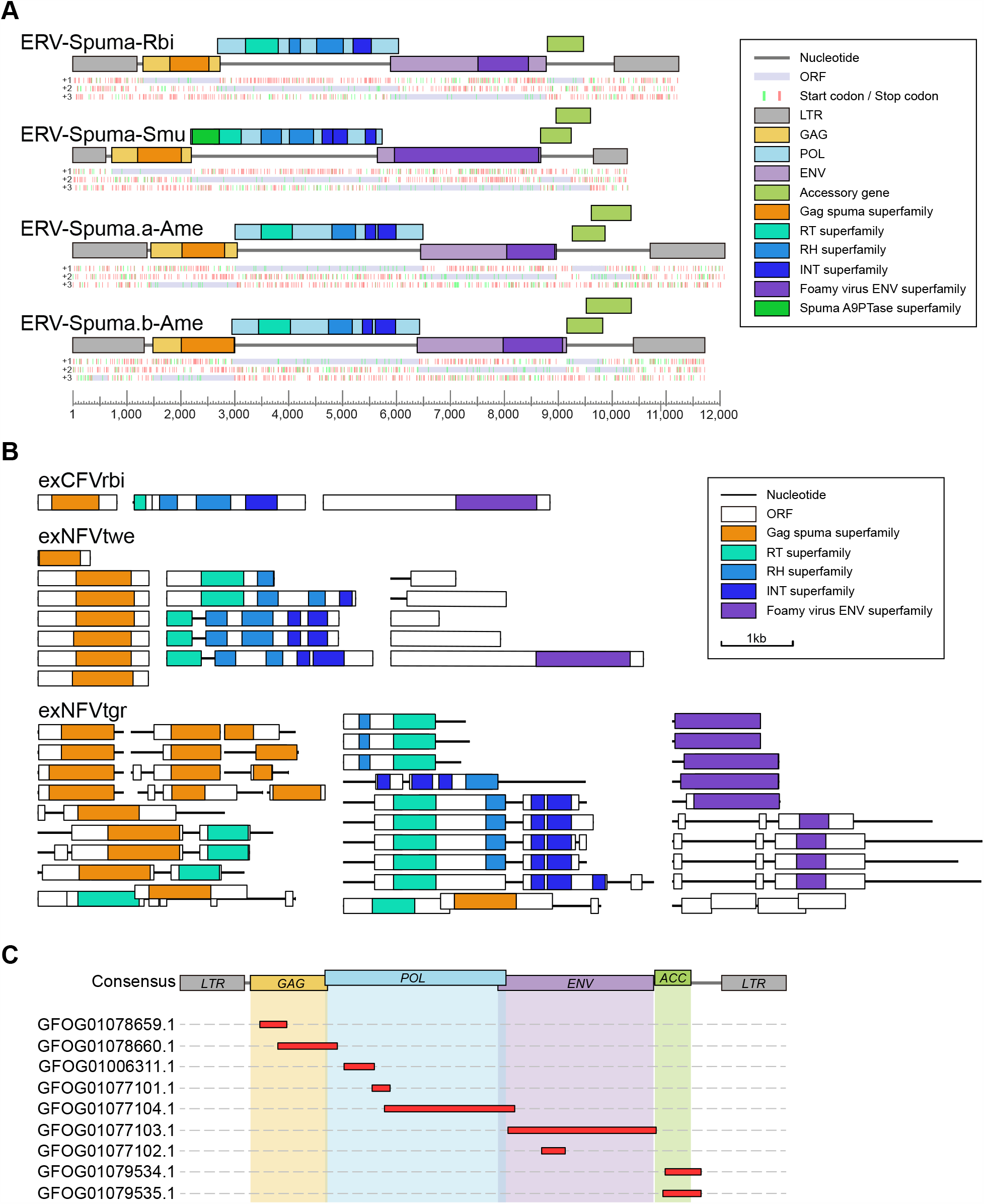
Genomic organization of novel EFVs and exFVs. **A)** The consensus genomic organizations of four novel EFVs. The consensus genomes of EFVs are drawn as lines and boxes to scales. The distribution of stop (red) and start (green) codons in three forward frames ((+1, +2, +3); from top to bottom) are shown under genomic schematic diagram of each consensus genome. Putative open reading frames (ORFs) are shown in light purple and were used to determine viral coding regions. The predicted domain or regions that encode conserved proteins are labelled with colored boxes. **B)** the representative genomic structures of exFVs. the contigs and ORFs of expFVs are drawn as lines and boxes to scale. The predicted domain or regions that encode conserved proteins are labelled with colored boxes. **C)** Mapping result of the exCFVrbi to consensus ERV-Spuma-Rbi genome. LTR, long terminal repeat; GAG, group-specific antigen gene; POL, polymerase gene; ENV, envelope gene. RT, reverse transcriptase; RH,;

The genomes of exFVs were then separately annotated and we found that all lineages of exFVs harbored major genes, including gag, pol and env, and they were distributed in different contigs in most cases (Fig. 2B). Accordingly, the foamy conserved domains were also identified in each lineage, including: (1) the Gag_spuma super family domain (cl26624)(40, 41) and the (2) Foamy_virus_ENV super family domain (cl04051)(38) and other retroviral domains (RT, RH and IN). We also found that most copies of exNFVtgr contained premature stop codons or only harbored partial genes, which indicated that they might be ERV-derived RNA. It is worth noting that exCFVrbi-1 (containing gag), exCFVrbi-2 (containing pol), exCFVrbi-3 (containing env), exNFVtwe -1,2,4,5 (containing gag), exNFVtwe-4 (containing pol), and exNFVtwe-5 (containing env) harbored major genes separately without any stop codon or indels, indicating the possible existence of complete functional genomes for exCFVrbi and exNFVtwe.

As both exCFVrbi and EFV have been found in *R. bivittatum*. We then further checked the homology of these two viruses (Fig. 2C). By mapping the assembled exFV contigs to a consensus ERV-Spuma-Rbi, we found that all the exFV contigs showed high (98% - 100%) similarity to the latter, indicating they were the same foamy virus. Surprisingly, we found that the exCFVrbi contigs were not identical to any copies of ERVs-Spuma-Rbi, and combined with the fact that exCFVrbi contained all major proteins (gag, pol, env and accessory gene) without any stop codons or indels, these results indicated the high probability of the existence of exogenous foamy viruses in *R. bivittatum*.

### Phylogenetic analysis

To elucidate the relationship between novel exFVs and EFVs identified here with other vertebrate FVs and EFVs, both long Pol (> 500 amino acid residues in length) and Env (> 320 aa in length) protein phylogenetic trees were generated to accommodate for their different evolutionary histories (Fig. 3). The phylogeny of pol and env were slightly different, supporting the theory that different genes of FVs indeed had different evolutionary histories(32, 37, 42). However, both phylogenies indicated that the six lineages of EFVs discovered in amphibians could be divided into three clades, giving support to the idea that amphibian FVs had multiple origins. Two lineages of ERV-Spuma-Ame and two exNFVs found in salamander clustered together with the previously identified NviFLERV, which was consistent with our co-divergence theory. Also, ERV-Spuma.a-Ame and ERV-Spuma.b-Ame robustly clustered together with a long branch, re-confirming that they independently resulted from two different foamy virus infections. In addition, in the pol phylogeny, the novel *Gymnophiona* exCFVrbi and ERV-Spuma-Rbi sequences formed a sister clade closely related to salamander FVs, and they formed a monophyletic group with robust support in the env phylogeny, indicating the different evolutionary histories of genes from *Gymnophiona* FVs. Notably, frog ERV-Spuma-Smu formed a well-supported monophyletic group that was most closely related to avian EFVs, indicating that ERVs-Spuma-Smu were possible acquired from cross-species transmission rather than through virus-host divergence.

**Fig. 3.**
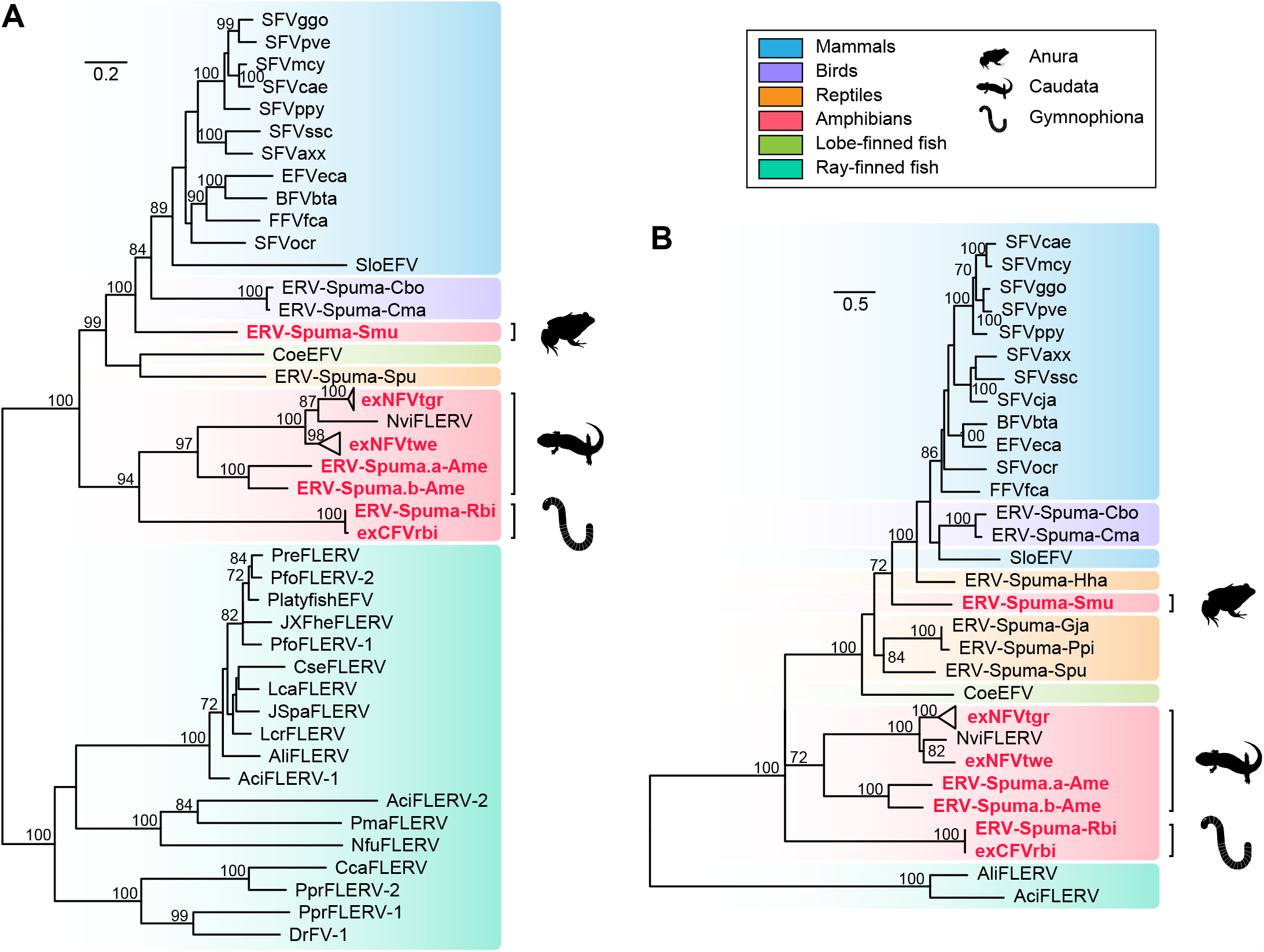
Phylogenetic trees of FVs, exFVs and EFVs. The trees were inferred using amino acid sequences of the **A)** Pol gene and **B)** Env gene. The trees are midpoint rooted tree for clarity only. The newly identified exFVs and EFVs are labeled in red. The scale bar indicates the number of amino acid changes per site. Bootstrap values <70 percent are not shown.

### Relationship of foamy viruses with their hosts

Previous research has provided strong evidence for the co-divergence of foamy viruses and their hosts, and some cross-species transmission events have also been observed in EFVs(18, 29, 34-36). Here, to further investigate the deep histories and evolution relationships between FVs and their vertebrate hosts, we generated a phylogenetic tree for FVs, EFVs and exFVs (Fig. 4). This tree showed that most FVs maintained a stable co-divergence pattern with their hosts (Fig. 4A and 4B). However, cross-species transmission also played an important role in the early evolution of foamy viruses, specifically in amphibians, where frog ERV-Spuma-Smu formed a single clade and was close related to avian EFVs rather than other amphibian FVs. In addition, exNFVtgr clustered with exNFVtwe, which was also inconsistent with their host phylogeny. We also noted that *Gymnophiona* ERV-Spuma-Rbi and exCFVrbi formed a well-supported (bootstrap = 88) monophyletic clade, giving additional credence to the idea that amphibian FVs had multiple origins and contained several paraphyletic groups.

**Fig. 4.**
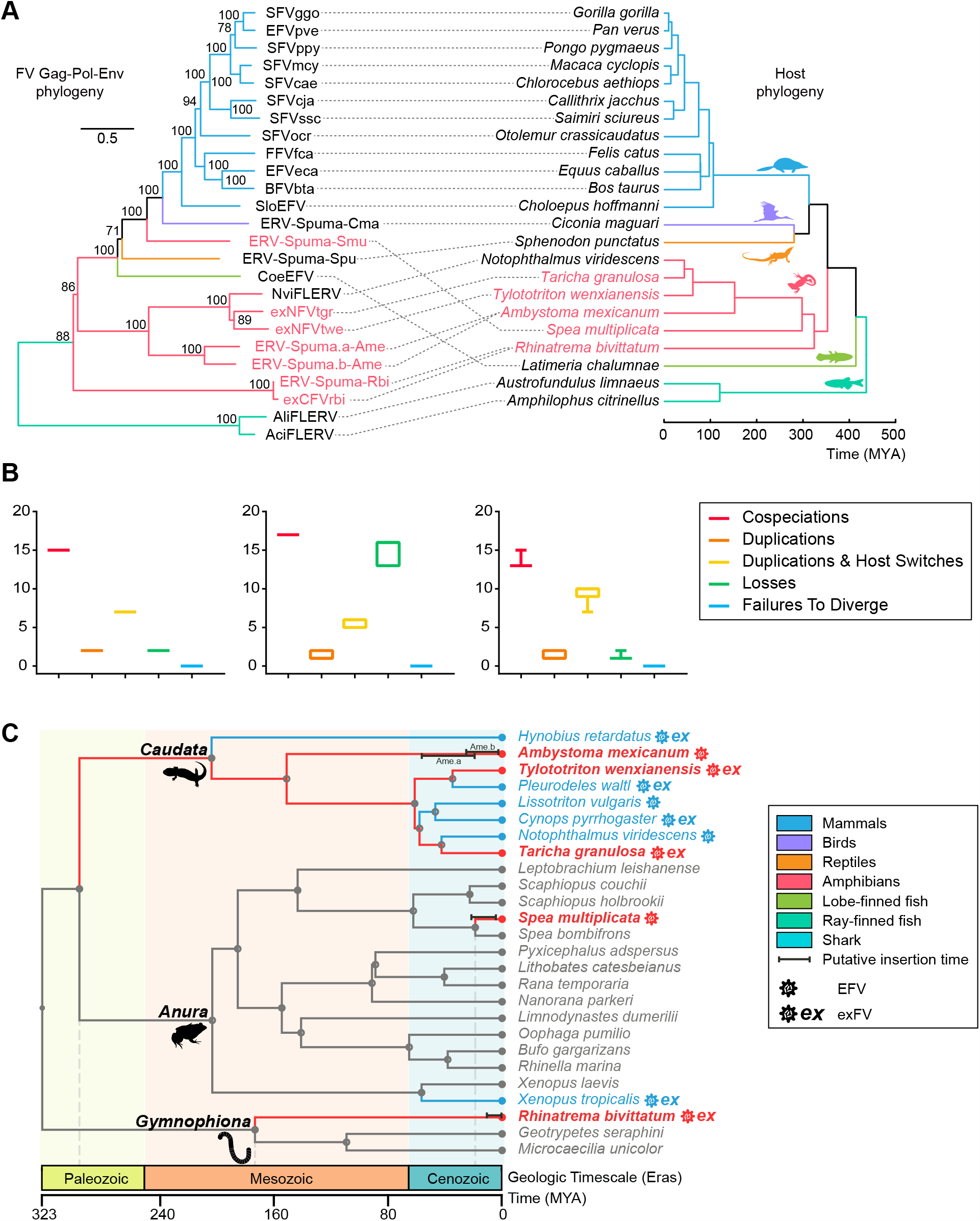
The macro-evolutionary history of foamy viruses and their vertebrate hosts. **A)** Association between foamy viruses (left) and their hosts (right). Associations between foamy viruses and their hosts are indicated by connecting lines. The newly identified exFVs, EFVs and their hosts are labeled in red. Scale bars indicate the number of amino acid changes per site in the viruses and the host divergence times (million years ago, MYA). Bootstrap values <70 percent are not shown. **B)** reconciliation analysis of foamy virus using Jane. co-speciations (red), duplications (orange), host-switching (yellow), losses (green) and failures to diverge (blue) events. The cost parameters (cospeciation_duplication_duplication& host switch_loss_failure to diverge_failures to diverge) used in each test are left (-1_0_0_0_0), middle (0_1_2_1_1) and right (0_1_1_2_0). **C)** The calibration of foamy virus infection timeline. A time-calibrated phylogeny of screened amphibians is obtained from TimeTree (http://www.timetree.org/). Previous estimated endogenization intervals are labeled with blue lines while our predictions are labeled with red lines. Closed circles on nodes represent the existence of taxon rank names. The time range of LTR dating estimation for each foamy virus is shown with black bar.

To roughly estimate the insertion time of amphibian EFVs, an LTR divergence-based dating method was used(43, 44). In total, 3 ERVs-Spuma-Smu, 43 ERVs-Spuma.n-Rbi, 5 ERVs-Spuma.a-Ame and 3 ERVs-Spuma.b-Ame were included in our dating estimation (Table S4 and Fig. 4C). This analysis revealed that ERVs-Spuma-Smu were relatively young, and their insertions could be dated back to 3.7–20.9 million years ago (MYA), close to the estimated divergence time of *S. multiplicate* (18.9 MYA). However, the insertion of ERVs-Spuma-Rbi could be traced back to as recent as 12.2 MYA, which was much younger than the estimated divergence time for *R. bivittatum* (135–213 MYA). The insertion of ERVs-Spuma.b-Ame could be dated back to 2.4–24.9 MYA. On the contrary, another lineage in *A. mexicanum*, ERVs-Spuma.a-Ame, could be traced back to 18.9–56 MYA, which was more ancient. Nevertheless, as LTR dating might severely underestimate ERV ages, these estimates should be treated with caution(45).

Combining our co-evolution analyses with our dating estimations allowed us to further calibrate foamy virus infection timeline in amphibians (Fig. 4C). Including previous research, it seemed that most Salamander EFVs followed a co-divergent pattern (Fig.4a). To date, all screened salamander species harbored foamy viruses, 5/8 of which were exFVs, indicating the possible of circulation of foamy virus in Caudates which could date back to Paleozoic era and they together with their host appeared to have an ancient origin. On the contrary, frog EFVs and exFVs (ERV-Spuma-Smu and XtrFLERV) were appeared to be the result of cross-species transmission events (Fig. 4), and other species in the same genus of their host did not harbor such FVs(32). Thus, they were relatively young compare to other amphibian FVs, which emerged in the Cenozoic. However, this was the first time we found *Gymnophiona* EFV and exFV, and other related species did not harbor such FVs. Thus, their infection could date back to somewhere between the Mesozoic and Cenozoic era. Taking these facts into consideration, it seemed that origin of amphibian foamy viruses began somewhere between the Paleozoic and Cenozoic era.

## Discussion

In this study, we reported a distinct lineage of FVs in two-lined caecilian (order: *Gymnophiona*) at both the DNA and RNA level, which added *Gymnophiona* to the list of currently known hosts of FVs. To date, in amphibians, 8 salamanders, 2 frogs and 1 caecilian have been confirmed to carry FVs (Fig. 4A)(32). However, the evolution histories of FVs in different hosts appeared to be extremely different (Fig. 4). We found that all screened salamanders harbored FVs and they maintained a co-divergence pattern with their host within the clade of salamanders(32). It seemed that FVs might circulate in salamanders starting from the Paleozoic (283–311 MYA). On the contrary, only 2 of 15 frogs carried FVs, indicating the rarity of FVs in frogs, and they were largely not a reservoir for FVs. Notably, ERV-Spuma-Smu and the previously identified XtrFLERV in frogs were acquired from cross-class transmission recently in the Cenozoic era, where the former was from a potential unknown land retrovirus and the latter was from a marine retrovirus(32). This re-confirmed that the water-land interface was not a strict barrier to viral transmission and cross-class transmission occurred occasionally(9). In caecilians, although only 3 genomes were available, we were able to identify a lineage in two-lined caecilian. Phylogenetic analysis revealed that ERV-Spuma-Rbi was basal to all amphibians FVs, indicating the ancient origin of caecilian FVs (Fig. 4B). Taking all these into consideration, it seemed that amphibian FVs had multiple origins and complex evolutionary histories. However, it is possible that this pattern will change with a larger sampling of taxa such that the EFV phylogeny expands(34).

Previous research identified one salamander (Japanese fire belly newt) and four fishes that harbored two EFV lineages(32). Here, we identified a novel salamander (axolotl), which harbored multiple lineages of FVs and they were characterized at the genomic level. In fact, we found three lineages of FVs in axolotl (Fig. 2 and S3), including ERV-Spuma.a-Ame, ERV-Spuma.b-Ame and ERV-Spuma.c-Ame. However, we only identified one full-length ERV-Spuma-c-Ame among multiple copies (9 copies). Accordingly, the genome structure of this full-length ERV-Spuma.c9-Ame is presented in Fig. S3, but its predicted pairwise LTR was too short (215 bp) in length and it only harbored a partially conserved domain (GAG, RT and IN), which made it relatively difficult to align with other EFVs lineages. Thus, we did not include this lineage in our major analyses. However, taking this lineage into consideration, we confirmed that amphibians could be infected with several different FVs. These FVs, however, were more likely to integrate in host genomes in different periods of time (Fig. 4A). By comparing their genomes, we found that their envelope proteins showed limited similarity (46%). As the envelope gene determines the host range and binding receptor for a virus(46-49), this observation suggested that these two lineages might bind to different receptors when they infected their host.

Transcriptome data provided us with additional resources for discovering novel exogenous and endogenous viruses(50-52). Previous research has identified an enormous amount of viruses that have shown limited similarity to well-defined viruses(50, 53-55) and foamy viral contigs were also discovered in several salamanders (Fig. 4A)(26, 32). In our study, we improved the method by incorporating the method in virome analyses. This led to the discovery of three lineages of exFVs in two salamanders and one caecilian, and these viruses showed limited similarity to known exogenous mammal FVs (26%–39% conservation of the pol protein). We characterized these viruses in detail and found that all lineages harbored major proteins of retroviruses in different segments (Fig. 2B), which could not be directly assembled at the genome level. Overall, the efficacy of virus discovery using transcriptome analysis depends on sequencing depth and assembly quality. However, the workflow presented here provides a new refinement for discovering novel retroviruses.

Importantly, among all exFVs, we found that exCFVrbi contained no stop codons or any indel in our contigs, which included complete major genes (Fig. 2B). As both genome and transcriptome data were available for *R. bivittatum*, we made a comparison between exCFVrbi and ERV-Spuma-Rbi, and found that exCFVrbi was not identical to any copies of EFV. Thus, exCFVrbi could have been the first exogenous foamy virus in amphibians, although, we could not directly examine such viral particles. This research still supports the assumption that exogenous foamy viruses could exist in other species beside mammals. Nevertheless, it is also possible that errors in sequencing or assembly may have led to such phenomenon.

Recently, another study identified the exaptation of foamy virus in gecko(36). This indicated that foamy virus-derived genes could also be co-opted. In our study, we annotated exFV contigs in detail and found that 7 contigs contained gag and 3 contigs contained env in NFVtwe with no stop codon, which all had the capacity to be translated into proteins. In other words, they had potential as retroviral-derived candidates for co-option identification. Also worth noting was that most exFVtgrs contained stop codons and their ORFs were incomplete, which might have indicated that they were likely to be ERV-derived long noncoding RNAs (lncRNAs) rather than exogenous retroviruses. Moreover, they might also function as lncRNAs to participate in genomic regulatory processes(56-58). However, this speculation should be verified by further study.

In conclusion, by integrating genomics and transcriptomic data and performing a phylogenomic analysis, we discovered 6 lineages of FVs that had different evolutionary histories, doubling the known set of foamy viruses in amphibians. We also confirmed that amphibians could be infected by multiple FVs at different periods of time. Interestingly, we identified the first potential exogenous form of FV (exCFVrbi) circulating in caecilian, which supports the idea that exogenous foamy virus could exist in other species beside mammals. This research demonstrated that repeated infections and multiple origins for amphibian FV sand revealed a complex macroevolution of foamy viruses with their hosts.

## Materials and Methods

### Genome and transcriptome screening and EFV/exFV identification

As most EFVs showed limited similarity to exogenous FVs and previously found EFVs, a stepwise method was used for amphibian EFV mining. First, all 19 amphibian genomes (Table S1) in Genbank as of November 2020 were screened for foamy-like viruses using tblastn(59), and conserved Pol proteins of foamy viruses, including EFVs, were used as probes (Table S3). A 25% sequence identity over a 40% region with an e-value set to 1E-5 was used to filter significant hits. Second, potential foamy-like elements were included in phylogenetic analysis. Hits that clustered with EFVs and FVs were considered EFVs. Then, the flanking sequences of these EFVs were extended to identify viral pairwise LTRs using BLASTN(59), LTR_Finder and LTR_harvest. In total, we were able to identify 5 full-length (containing a pairwise LTRs) EFVs in *S. multiplicate*, 46 in *R. bivittatum*, and 26 in *A. mexicanum*. These full-length EFVs were then used as a query to search for EFV copies using blastn. Sequences longer than 4 kb with 85% identity were regarded as copies of each EFV lineage (Table S2). In accordance with the nomenclature proposed for ERVs, EFVs found in *S. multiplicate, R. bivittatum*, and *A. mexicanum* were named ERVs-Spuma.n-Smu, ERVs-Spuma.n-Rbi, and ERVs-Spuma.n-Ame, respectively. As there were 3 lineages of EFVs in *A. mexicanum*, they were separately designated as ERVs-Spuma.an-Ame, ERVs-Spuma.bn-Ame, and ERVs-Spuma.cn-Ame, respectively.

To identify potential exFVs, transcriptome sequencing assembly database (TSA) were screened using tblastn and all three major proteins of amphibian EFVs, including the newly identified ERVs-Spuma-Smu, ERVs-Spuma-Rbi and ERVs-Spuma-Ame were used as probes. A 25% sequence identity over a 40% region with an e-value set to 1E-5 was used to filter significant hits. Then, the calling hit contigs were included in the phylogenetic analysis. The viral contigs within a clade of FVs were considered. In total, three exFVs were found and exFVs in *T. granulosa, T*.*wenxianensis*, and *R. bivittatum* were named as exNFVtgr.n, exNFVtwe.n, and exCFVrbi.n, respectively.

### Consensus genome construction and genome annotation

EFVs longer than 5 kb in each EFV lineage were aligned using MAFFT 7.222(60) and then used to construct consensus sequences for each EFV lineage. The distributions of open reading frames (ORFs) in copies of EFV and exFV contigs were determined using ORFfinder (https://www.ncbi.nlm.nih.gov/orffinder/) at NCBI and confirmed by Blastp(59). Conserved domains for each sequence were found by using CD-Search against the Conserved Domain Database (CDD) (https://www.ncbi.nlm.nih.gov/cdd/)(61).

To construct putative full-length coding regions for exFVs, we used the method shown below(51, 62). If a contig that mapped to the same coding gene (e.g., the pol gene) with high similarity (> 85%), then these contigs were used to construct consensus sequences for each gene. Otherwise, to decide which contig was used to construct genomes for exFVs, we: 1) compared the phylogenetic positions of related viral proteins; 2) compared the similarity with related proteins from a reference foamy virus; and 3) checked the completeness of our conserved domain. Then, the selected contigs were used to construct putative genomes for exFVs. However, the putative genomes for exFVs were only used in concatenated Gag-Pol-Env phylogeny.

### Molecular dating

ERV integration time can be approximately estimated using the relationship: T = (D/R)/2, in which T is the integration time (million years, MY), D is the number of nucleotide differences per site between a set of pairwise LTRs, and R is the genomic substitution rate (nucleotide substitutions per site, per year)(43, 44). We used a previously estimated neutral nucleotide substitution rate for frogs (9.24 × 10^−10^ to 1.53 × 10^−9^ nucleotide substitutions per site, per year)(63) to estimate the evolution of amphibians. LTRs less than 300 bp in length were excluded in this analysis(35, 64). In total, 3 ERVs-Spuma -Smu, 43 ERVs-Spuma.n-Rbi, 5 ERVs-Spuma.a-Ame, and 3 ERVs-Spuma.b-Ame containing a pairwise intact LTR were used to estimate integration time in this manner (Table S4).

### Phylogenetic analysis

To investigate the evolutionary relationship between FVs, exFVs and EFVs, protein sequences for RT, Pol, Env and concatenated Gag-Pol-Env were aligned using MAFFT 7.222(60)(Data set S2-5). The regions in the alignment that aligned poorly were removed using TrimAL(65) and confirmed manually in MEGA X(66). A sequence was excluded if its length was less than 75% of the alignments. The best-fit models (RT: VT+G; Pol: LG+F+I+G; Env: LG+F+I+G; and Gag-Pol-Env: LG+F+I+G) were selected using ProtTest^49^ and the phylogenetic trees for these protein sequences were inferred using the maximum likelihood (ML) method in PhyML(67) or IQ-Tree(68) incorporating 100 bootstrap replicates to assess node robustness. Phylogenetic trees were viewed and annotated in FigTree V1.4.3 (https://github.com/rambaut/figtree/).

### Co-evolution analysis

To assess the macroevolution of foamy viruses and their hosts, event-based Jane 4(69) was used. We set the cost parameter (cospeciation, duplication, duplication& host switch, loss and failure to diverge), based on previous research, as 1) -1,0, 0, 0, 0(32); 2) 0, 1, 2, 1, 1; and 3) 0, 1, 1, 2, 0(69). Then Jane performed statistical analyses to assess robustness by generating random parasite trees with a sample size of 500.

## Data availability

All the data needed to generate the conclusions in the article are present in the article itself and the supplementary data.

## Acknowledgments

This work was supported by the National Natural Science Foundation of China (31671324 and 31970176) and CAS Pioneer Hundred Talents Program.

## Conflict of interest

None declared.

